# Characterisation of the transcriptome and proteome of SARS-CoV-2 using direct RNA sequencing and tandem mass spectrometry reveals evidence for a cell passage induced in-frame deletion in the spike glycoprotein that removes the furin-like cleavage site

**DOI:** 10.1101/2020.03.22.002204

**Authors:** Andrew D. Davidson, Maia Kavanagh Williamson, Sebastian Lewis, Deborah Shoemark, Miles W. Carroll, Kate Heesom, Maria Zambon, Joanna Ellis, Phillip A. Lewis, Julian A. Hiscox, David A. Matthews

## Abstract

Direct RNA sequencing using an Oxford Nanopore MinION characterised the transcriptome of SARS-CoV-2 grown in Vero E6 cells. This cell line is being widely used to propagate the novel coronavirus. The viral transcriptome was analysed using a recently developed ORF-centric pipeline. This revealed the pattern of viral transcripts, (i.e. subgenomic mRNAs), generally fitted the predicted replication and transcription model for coronaviruses. A 24 nt in-frame deletion was detected in subgenomic mRNAs encoding the spike (S) glycoprotein. This feature was identified in over half of the mapped transcripts and was predicted to remove a proposed furin cleavage site from the S glycoprotein. This motif directs cleavage of the S glycoprotein into functional subunits during virus entry or exit. Cleavage of the S glycoprotein can be a barrier to zoonotic coronavirus transmission and affect viral pathogenicity. Allied to this transcriptome analysis, tandem mass spectrometry was used to identify over 500 viral peptides and 44 phosphopeptides, covering almost all of the proteins predicted to be encoded by the SARS-CoV-2 genome, including peptides unique to the deleted variant of the S glycoprotein. Detection of an apparently viable deletion in the furin cleavage site of the S glycoprotein reinforces the point that this and other regions of SARS-CoV-2 proteins may readily mutate. This is of clear significance given the interest in the S glycoprotein as a potential vaccine target and the observation that the furin cleavage site likely contributes strongly to the pathogenesis and zoonosis of this virus. The viral genome sequence should be carefully monitored during the growth of viral stocks for research, animal challenge models and, potentially, in clinical samples. Such variations may result in different levels of virulence, morbidity and mortality.

## Introduction

Since the emergence of severe acute respiratory syndrome coronavirus-2 (SARS-CoV-2) as a human pathogen at the end of 2019, the virus has spread globally, causing > 312 000 confirmed cases of COVID-19 and 13,400 deaths as of the 22^nd^ March, 2020^1^. Although vaccines are under rapid development to prevent SARS-CoV-2 infection, little is known of either the immune correlates of protection or the ability of the virus to avoid the host immune response through mutation and recombination^2^.

The genome sequence was rapidly determined and revealed that SARS-CoV-2 is most closely related to bat coronavirus RaTG13 and clusters with SARS-CoV, that emerged in 2002, in the genus *Betacoronavirus* of the family *Coronaviridae*^3,4^. Based on homology to other known coronaviruses^3,4^ the genome sequence was used for viral transcript prediction and the annotation of ORFs. Coronaviruses use a complex strategy to express their genetic information^5^, and transcription is initiated from defined (and sometimes cryptic) transcription associated sequences^6^. Similar, to other coronaviruses, the SARS-CoV-2 29.9 kB RNA genome contains two large ORFs, ORF1a and ORF1ab, predicted to be initially translated into the polyproteins, pp1a, and pp1ab that arises by ribosomal frameshifting (Figure 1A). The polyproteins are post-translationally processed by viral encoded proteases to produce 16 proteins that are conserved between coronaviruses and proposed to function in the synthesis of viral RNA and immune evasion^7^. During viral genome replication a set of “nested” subgenomic mRNAs are produced that are predicted to encode the structural proteins spike (S), envelope (E), membrane (M) and nucleocapsid (N) and at least nine small accessory proteins, some of which are unique to SARS-CoV-2^3,4^. The subgenomic mRNAs have a common 5’ leader sequence and are 3’ co-terminal with a polyA tail. This feature is likely to be added by viral encoded proteins in the cytoplasm of an infected cell.

**Figure 1.**
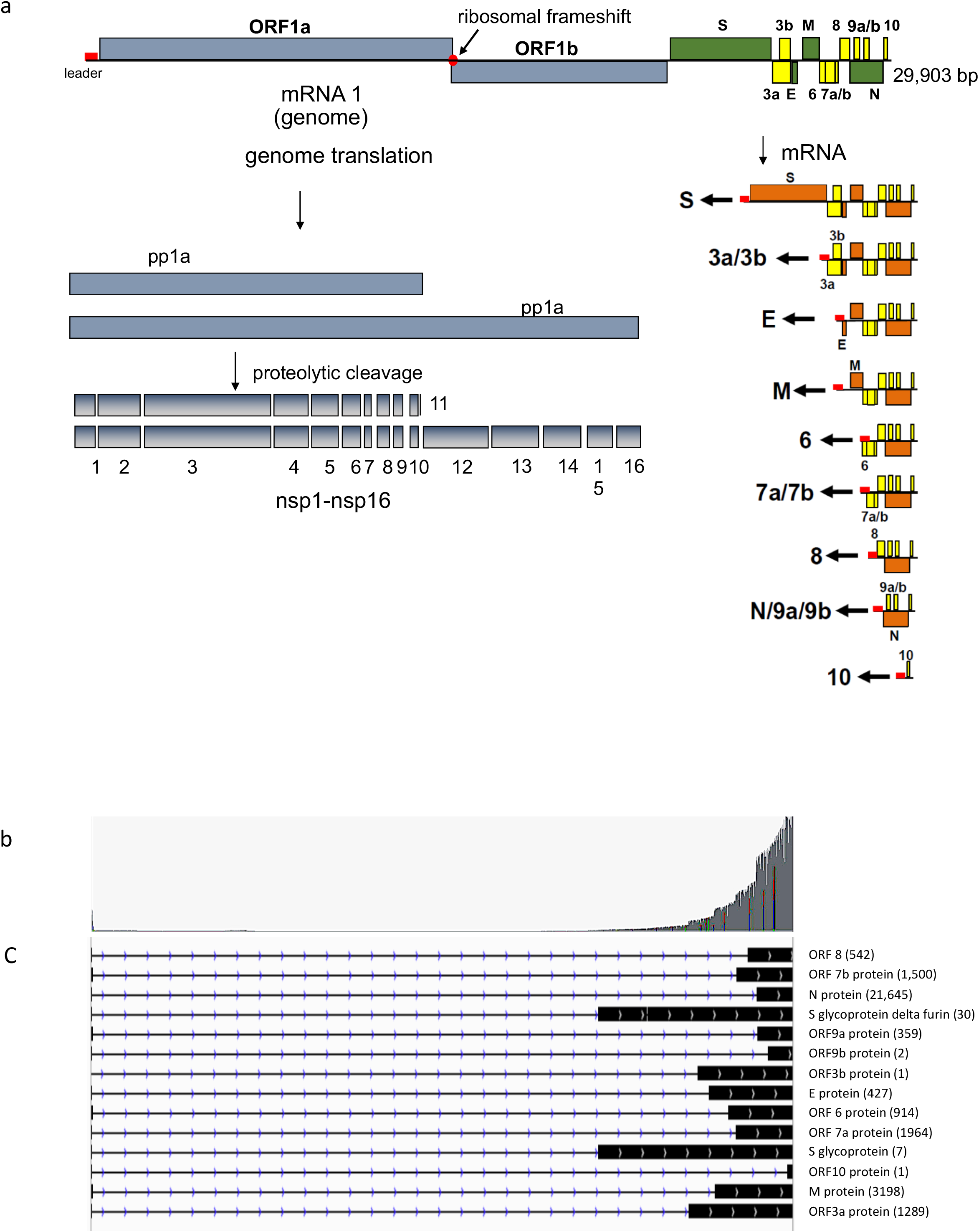
Overview of nanopore inferred transcriptome. Part a illustrates the classical transcription map of coronaviruses adapted for SARS-CoV-2. The genome is itself an mRNA which when translated gives rise to polyproteins pp1a and, upon a ribosomal frameshift, pp1ab. These polyproteins are proteolytically processed down to a range of non-structural proteins termed nsp1-16 some of which will form the viral replication-transcription complex (RTC). The RTC then generates subgenomic mRNA which canonically contains a sequence present at the 5’ end of the viral genome known as the Transcription Regulating Sequence - Leader (TRS - L) linked by discontinuous transcription to one of several similar sequences downstream of pp1ab. In this manner the remaining ORFS on the viral genome are placed 5’ most on the subgenomic mRNA and are subsequently translated. Part b shows the total read depth across the viral genome for all reads, the maximum read depth was 511,129. Part c illustrates the structure of dominant transcript coding for each of the identified ORFS. Only transcripts that starts inside the leader TRS sequence are considered here. The black rectangles represent mapped nucleotides and the arrowed lines represent regions of the genome that are skipped. To the right is noted the 5’ most ORF is that the transcript would code for, in parenthesis we note how many individual transcripts were observed.

Recently, the SARS-CoV-2 transcriptome has been examined using direct RNAseq (dRNAseq)^8^,^9^. Furthermore, a recent proteomic study showed that a number of the predicted viral proteins are produced in infected cells, but not all of the predicted viral proteins were detected^10^. We pioneered the use of combining transcriptomics and proteomics to study viral infections *in vitro* and *in vivo*, initially using the well characterised respiratory virus, adenovirus but also respiratory syncytial virus and highly pathogenic zoonotic viruses such as Hendra virus^(11–13^. More recently we have focussed on the latest technologies to study the transcriptome of viruses, including direct RNA sequencing on nanopore devices using long read length sequencing. In particular, we developed a novel ORF-centric pipeline to analyse the very large amounts of transcriptomic data generated by this technique^14^. This pipeline was first used on data generated from adenovirus infected cells. The approach rapidly reproduced transcript maps that correlated highly with the previously described transcriptome maps of adenovirus which have been carefully curated over several decades.

The SARS-CoV-2 subgenomic mRNAs encoding the structural proteins and their expression profiles are of particular interest as the encoded proteins are likely targets of a protective immune response. The coronavirus S glycoprotein is present as a homotrimer, protruding from the surface of the virion. It is a key determinant of viral tropism and the major antigenic target^(15,16^. The S glycoprotein is a class I fusion protein comprised of two domains, S1 and S2, responsible for receptor binding and the fusion of viral and cellular membranes respectively^(17,18^. The coronavirus S glycoprotein of coronaviruses is primed for cell entry by host cell proteases that cleave the protein at one or more positions. Cleavage of a “S1/S2” site at the boundary of the S1 and S2 domains occurs for some coronaviruses whilst cleavage at a “S2” site is common to all coronaviruses and results in activation of a highly conserved fusion peptide immediately downstream of the S2 site^(19–21^. Although the SARS-CoV-2 S glycoprotein shares 97% identity with that of bat coronavirus RaTG13^22^, an important difference is the presence of a four amino acid insertion (681PRRA684) at the S1/S2 cleavage site that introduces a potential furin-like cleavage site^(23,24^. The furin cleavage site is also not found in other “lineage B” betacoronaviruses including SARS-CoV, but has been detected in betacoronaviruses in other lineages, suggesting that this region has arisen during natural evolution^25^.

Here we used a combined transcriptomics and proteomics approach to produce a correlated transcriptome and proteome map of SARS-CoV-2 in the African Green monkey kidney cell line Vero E6. This cell line is routinely used globally to propagate viruses from clinical samples as well as to generate stocks of virus for academic research, drug susceptibility testing and vaccine challenge studies. There was a close correlation between predicted viral transcriptome and proteome maps as well as novel transcripts that encoded for proteins detected by tandem mass spectrometry. Finally, evidence was gathered for an in-frame deletion inside the S glycoprotein that removed the proposed furin cleavage site.

## Results

Complete characterisation of a novel and dangerous pathogen at a molecular level is highly desirable. Although analysis of the SARS-CoV-2 genome enables high confidence prediction of potential transcripts and ORFS, there is a pressing need to have the predictions confirmed and novel transcripts identified and assessed for biological significance and role in virulence and pathogenesis. At the same time, independent confirmation that viral transcripts were expressed was achieved by tandem mass spectrometry-based proteomics. This provided direct observation of the whole range of viral proteins and helped resolve ambiguities in the transcriptomic data. Finally, identification of phosphorylation sites provided an invaluable list of therapeutic targets based on kinase inhibitors. Using proteomics informed by transcriptomics (PIT) ^11^ on duplicate infected cells offered a rapid and high resolution approach to these key molecular virology questions.

### Overview of dRNAseq outputs

Long read length sequencing using an Oxford Nanopore MinION on polyA+ RNA from cells infected with SARS-CoV-2 was used to characterise viral RNA species. This technique has recently been used to study herpes viruses, coronaviruses and adenovirus transcriptomes ^14,26,27^. In total, 1,588,330 sequences were base called and passed QC, of those 527,401 were mapped to the BetaCoV/England/02/2020 genome using minimap2 (Figure 1b). In addition, an average polyA tail length for each transcript was generated using nanopolish and this information was added to the analysis pipeline. A strict cut off of a 20 nt minimum polyA length was employed and 386,903 transcripts passed this test. Accurately determining the 5’ end of direct

RNAseq transcripts has been shown to be problematic ^14,27,28^. The accepted model for coronavirus transcription (Figure 1a) proposes that all viral mRNAs have a common 5’ end ^29^ and the analysis herein was further restricted to those transcripts that met this criterion which reduced the total number of identified full length subgenomic mRNA molecules to 72,124. Figure 1c illustrates a transcription map of those transcripts that start at the expected location and have a known ORF as the 5’ most ORF. Transcripts that could express all of the predicted ORFs were present, including for the newly predicted ORF10 ^30^. However, only one transcript was detected and as previously suggested ^(8,9^ the status of this as a genuine transcript needs careful further investigation. Moreover, non-canonical TRS joining events were detected leading to, for example, apparently bone fide subgenomic transcripts coding for ORF7b. this was also recently reported by Kim et al. ^8^. The distribution of transcripts with a 5’ UTR consistent with the model of coronavirus mRNA expression that would also code for a predicted protein are shown in Table 1. Also reported are the average polyA tail lengths of each transcript group. These appeared to be shorter than most human eukaryotic transcripts but is consistent with recently reported findings for SARS-CoV-2 ^8^. Finally, the data from the analysis pipeline was tabulated and describes the structure of transcripts, how often they were observed, what features were present on each transcript and where the dominant transcription start sites and junctions are (Supplementary Tables 1-4).

**Table 1.**
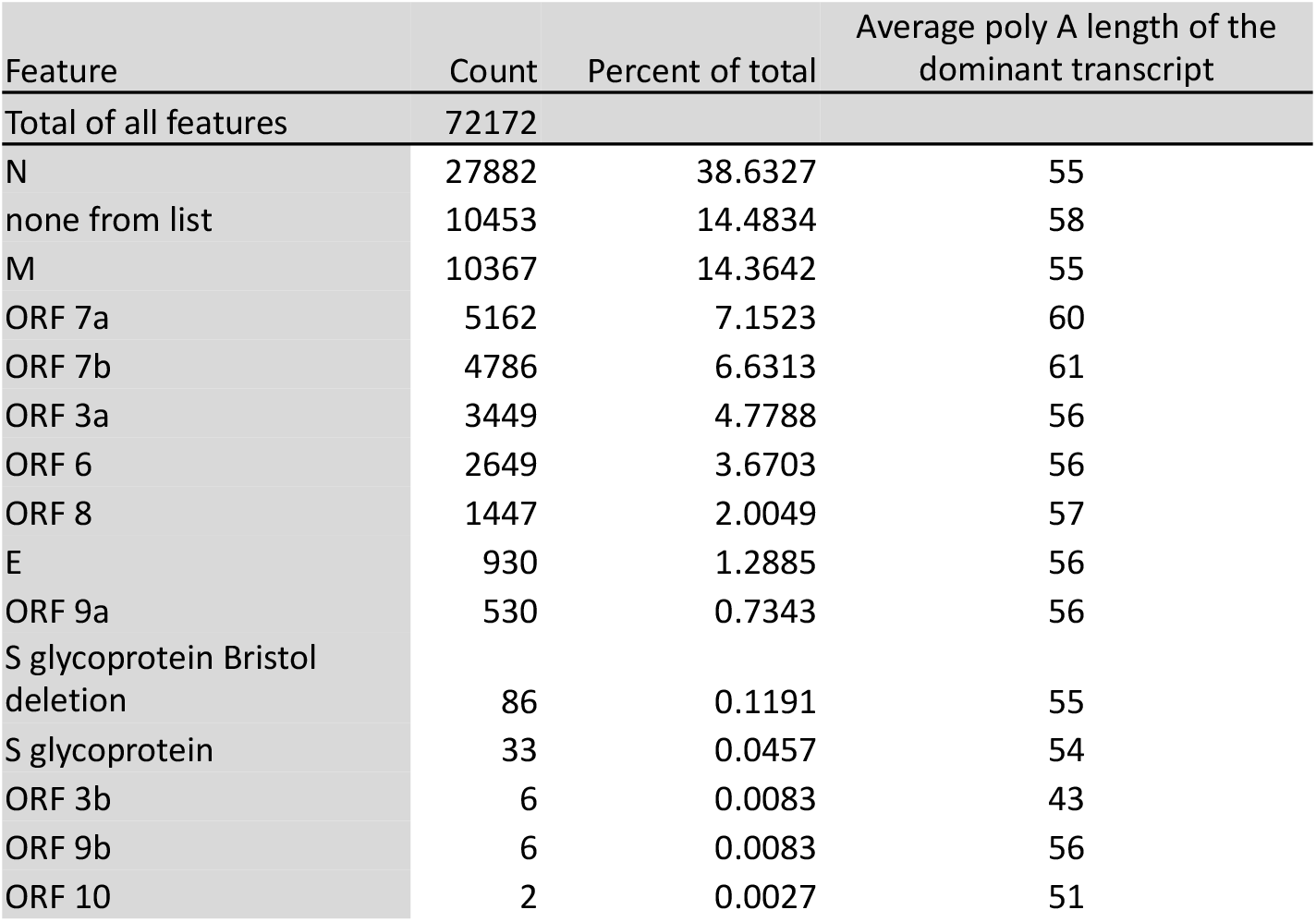
Count of transcripts where the 5’ most ORF is a recognised ORF. Only transcripts that start to map within the expected leader TRS are considered. For each transcript group the average polyA length is also shown for the dominant transcript that codes for the indicated ORF. Note that around 14% of transcripts do not apparently code for a recognised ORF, noted as “none from list”.

| Feature | Count | Percent of total | Average poly A length of the dominant transcript |
| --- | --- | --- | --- |
| Total of all features | 72172 |  |  |
| N | 27882 | 38.6327 | 55 |
| none from list | 10453 | 14.4834 | 58 |
| M | 10367 | 14.3642 | 55 |
| ORF 7a | 5162 | 7.1523 | 60 |
| ORF 7b | 4786 | 6.6313 | 61 |
| ORF 3a | 3449 | 4.7788 | 56 |
| ORF 6 | 2649 | 3.6703 | 56 |
| ORF 8 | 1447 | 2.0049 | 57 |
| E | 930 | 1.2885 | 56 |
| ORF 9a | 530 | 0.7343 | 56 |
| S glycoprotein Bristol deletion | 86 | 0.1191 | 55 |
| S glycoprotein | 33 | 0.0457 | 54 |
| ORF 3b | 6 | 0.0083 | 43 |
| ORF 9b | 6 | 0.0083 | 56 |
| ORF 10 | 2 | 0.0027 | 51 |

### Novel deletion in the S glycoprotein

Manual inspection of the transcripts aligned to the virus genome revealed a large number of reads that aligned to the gene encoding the S glycoprotein with a 24 nt deletion. This was predicted to result in a deletion of 9 aa, encompassing the proposed furin-like cleavage site (Figure 2a) with the replacement of a single isoleucine residue. Indeed the analysis revealed that there were a large number of transcripts that had both a polyA tail > 20 nt and a start at the proposed transcription start location, more than the full length S-glycoprotein (Table 1). To confirm this finding and examine if the deletion was present in the original stock samples at the PHE reference laboratory, the region was PCR amplified from the virus passaged at Bristol or from the original stock. These amplified fragments were sequenced using the Sanger method. This revealed that the Bristol stock was indeed a mixture of genomes but the original stock was solely wild type sequence. Subsequent examination of a very high quality dataset reported recently by Kim et al. ^8^ did not detect any evidence of similar significant deletions in the S glycoprotein in their data. However, their dataset contained a spontaneous deletion within the region of the genome coding for the E protein resulting in a 9aa in-frame deletion in almost two thirds of the mapped transcripts (Supplementary Figure 1).

**Figure 2.**
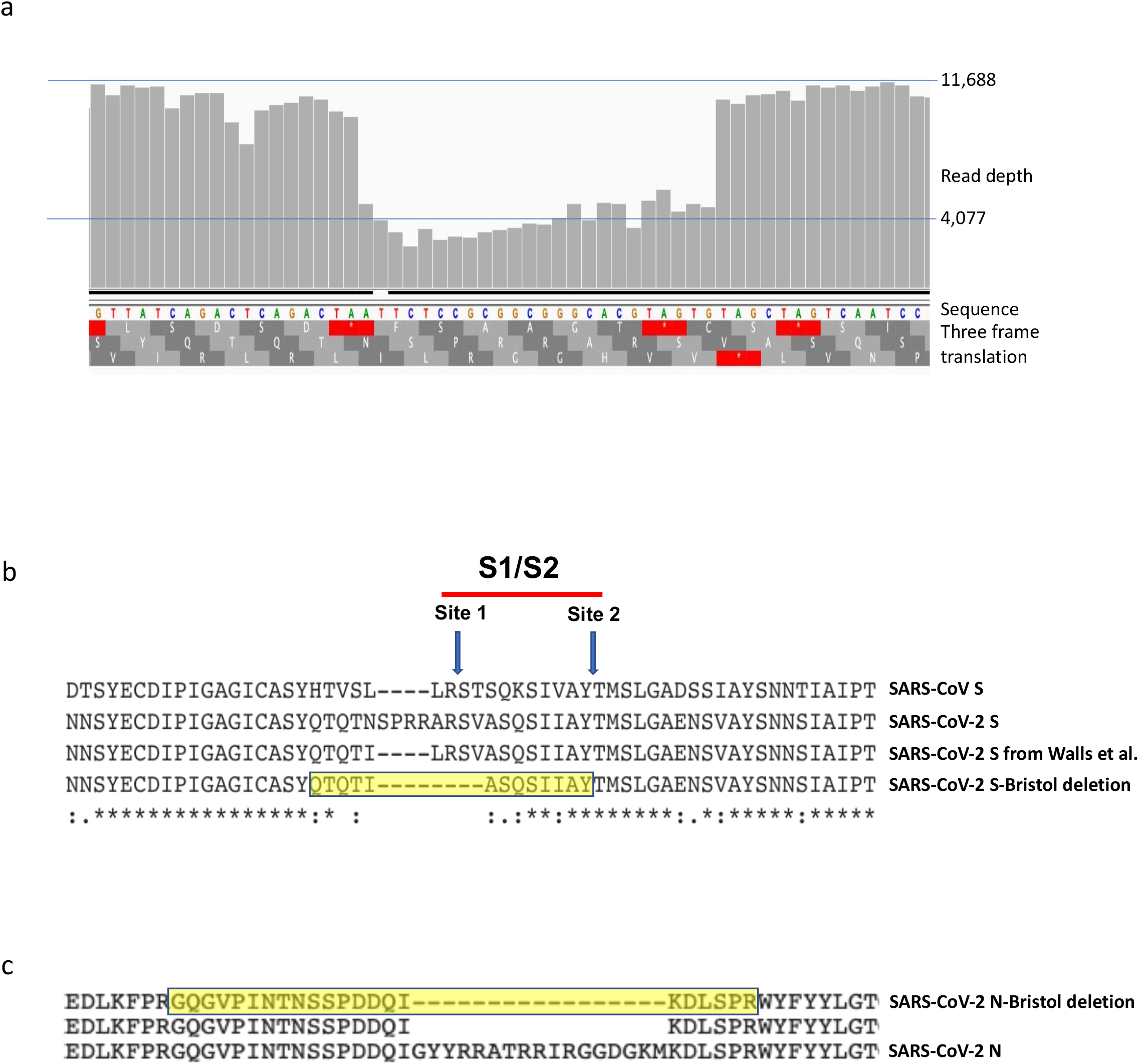
Deletions within the viral mRNA in the S glycoprotein and N In (a) is shown the read depth over the region deleted in the S glycoprotein together with information on the sequence in the region and the translation in all three frames. (b) is a clustal alignment of three proteins over this region, wild type SARS-CoV, wild type SARS-CoV-2, the artificially deleted version of the wild type SARS-Cov_2 S glycoprotein as reported in Walls et al. and finally the predicted sequence of the deleted protein described here. Highlighted in yellow is the sequence of the unique peptide generated by chymotrypsin digest of the protein which was identified by tandem mass spectrometry. (c) illustrates a similar examination of a proposed deletion in the N protein predicted by multiple aligned transcripts and subsequently identified in trypsin digested protein samples as indicated by the unique peptide highlighted in yellow.

### Proteomic detection of known SARS-CoV-2 proteins

Tandem mass spectrometry was utilised to detect as many viral peptides and phosphopeptides as possible (Supplementary dataset 5). Table 2 illustrates the numbers of peptides found for each of the predicted ORFs. Notably, whilst most of the ORFs detected were also confirmed in the recent report by Bojkova et al. ^10^, there were some differences. Whilst we could not detect peptides from ORF9b as described by Bojkova et al., peptides corresponding to the ORF9a protein were identified. This leaves only E, ORF7b and ORF10 to be detected by mass spectrometry. In addition, peptides unique to proteolytically processed components of the viral replicase polyprotein pp1ab; namely nsp4, nsp9, nsp12, nsp13, nsp14 and nsp15 were identified indicating that these important processing steps occurred as predicted (Table 3).

**Table 2.** Peptide counts for viral proteins. For each protein the total number of unique peptides is indicated alongside how many PSMs support the peptides identified. In the case of the viral polyprotein pp1ab, we also list how many peptides uniquely mapped to each nsp region.

| Protein Name | Unique peptides | PSMs | Polyprotein 1ab component | Unique peptides | PSMs | Polyprotein 1ab component | Unique peptides | PSMs |
| --- | --- | --- | --- | --- | --- | --- | --- | --- |
| N | 70 | 4152 | nsp1 | 10 | 59 | nsp10 | 5 | 18 |
| S Glycoprotein | 76 | 1984 | nsp2 | 41 | 277 | nsp12 | 40 | 146 |
| polyprotein 1ab | 323 | 1816 | nsp3 | 105 | 850 | nsp13 | 24 | 70 |
| M | 16 | 250 | nsp4 | 19 | 105 | nsp14 | 18 | 34 |
| ORF 3a | 8 | 119 | nsp5 | 9 | 54 | nsp15 | 20 | 50 |
| ORF 9a | 14 | 86 | nsp6 | 5 | 20 | nsp16 | 13 | 37 |
| ORF 8 | 3 | 33 | nsp7 | 4 | 13 |  |  |  |
| ORF 7a | 4 | 23 | nsp8 | 8 | 61 |  |  |  |
| ORF 6 | 2 | 3 | nsp9 | 4 | 20 |  |  |  |

**Table 3.** Peptides unique to processed proteins. The polyprotein pp1ab is processed into matured smaller proteins nsp1-16 during infection, this table indicates which unique peptides were identified that could only arise as a result of full polyprotein processing.

| Protein | Contributing PSMs | Sequence identified |
| --- | --- | --- |
| SARS2_pp1ab_nsp4 | 2 | ALNDFSNSGSDVLYQPPQTSITSAVLQ |
| SARS2_pp1ab_nsp9 | 1 | NNELSPVALR |
| SARS2_pp1ab__nsp12 | 4 | SADAQSFLNR |
| SARS2_pp1ab__nsp12 | 1 | YWEPEFYEAMYTPHTVLQ |
| SARS2_pp1ab__nsp13 | 2 | AVGACVLCNSQTSLR |
| SARS2_pp1ab__nsp14 | 2 | AENVTGLFK |
| SARS2_pp1ab__nsp15 | 1 | SLENVAFNVVNK |

Initially no unique peptides corresponding to the novel deleted version of the S-glycoprotein were detected by tandem mass spectrometry. However, a typical tryptic digest protocol would generate very large potentially undetectable peptides. Accordingly, samples were digested using chymotrypsin and the analysis repeated revealing a single peptide corresponding to the deleted version of the S-glycoprotein, but only when using a 5% false discovery threshold. However, this provided sufficient information to enable a targeted search. This identified nine Peptide Spectral Match’s (PSMs) with high confidence (i.e. FDR 1%) that corroborated the expression of the deleted spike protein (Supplementary dataset 6).

### Proteomic detection of previously unknown viral proteins

Transcriptomics and proteomics can be combined to provide unbiased insight into the proteome of viruses and higher eukaryotes ^11,12,31^. The ORF centric pipeline was used to generate a list of possible ORFs which could be made by translating the first ORF present on every transcript sequenced. As before ^14^, to overcome the inherent error prone nature of nanopore sequencing, the transcript was mapped to the viral genome and used the mapping information to create pseudo-transcripts based on the sequence of the viral genome.

Using this list of predicted proteins revealed a large number of novel peptides arising which could not be explained by the standard list of predicted viral proteins. In the majority of cases they appeared to be transcripts with rare and unusual structures likely resulting from rare rearrangements of the viral genome or potentially during subgenomic mRNA synthesis. However, in some cases a large number of similar transcripts were supported by direct peptide evidence that these rearrangements are indeed translated. In particular multiple versions of the N protein with distinct small internal deletions were detected for which 20 transcripts that could encode for these proteins were also identified (Supplementary dataset 7). A schematic of such a deletion in the N protein is shown, indicating the peptide that was identified by tandem mass spectrometry to support the observed transcripts (Figure 2c).

### Phophoproteomic analysis of SARS-CoV-2 proteins

Allied to an analysis of viral peptides, the phosphorylation status of viral proteins was investigated as this could reveal potential targets for licenced kinase inhibitors. Phosphopeptides corresponding to locations on the N, M, ORF 3a, nsp3, nsp9, nsp12 and S-glycoprotein were detected (Table 4). The presence of phosphorylation sites on the S glycoprotein has not been previously noted and may be of significance in terms of vaccines based on this protein, the location of these sites is illustrated (Figure 3). The location of the phosphorlaytion sites on the proteins listed is illustrated in Figure 4 and additional models of the phosphorylation sites on the N-terminal RNA binding domain of the N protein are shown in Figure 5.

**Table 4.**
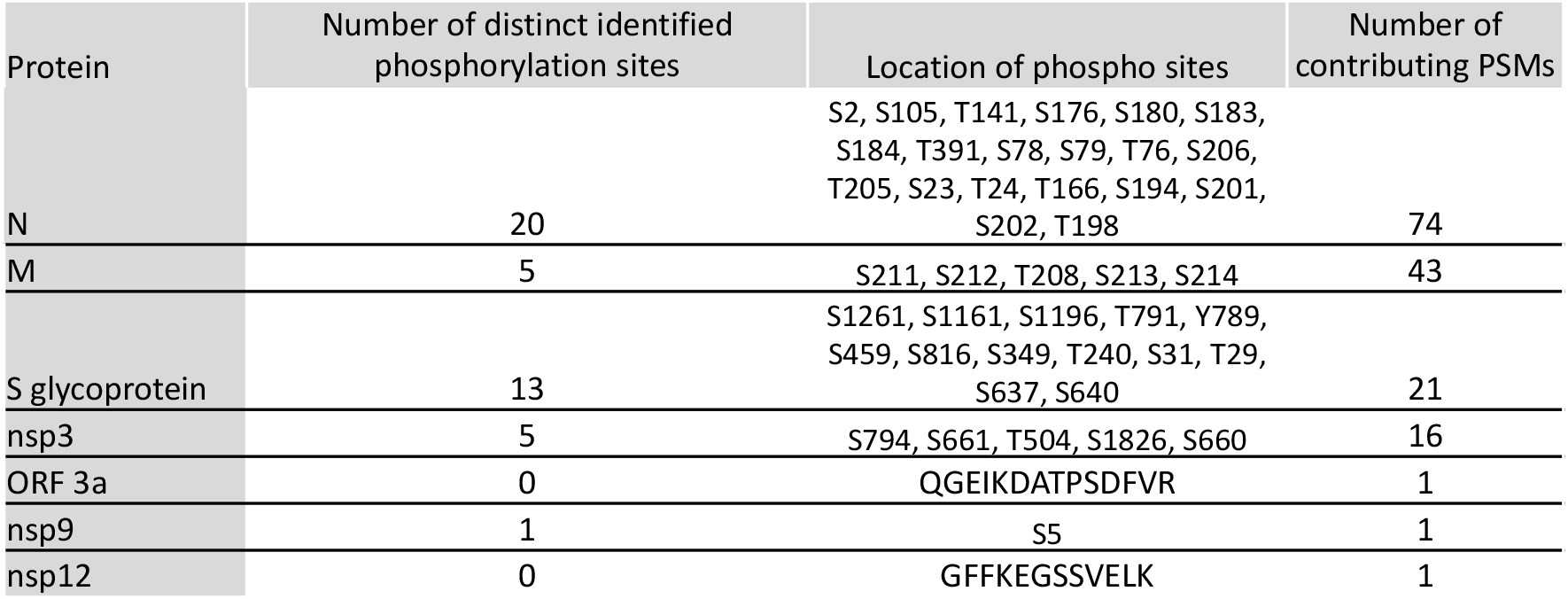
Phosphopeptide counts. Listed for each protein is the number of distinct phosphor sites identified along with the locations and amino acid modified as well as how many contributing PSMs there are in total. For proteins ORF3a and nsp12 no distinct site could be identified despite a phosphorylated peptide being found, in these cases the peptide sequence is provided.

**Figure 3.**
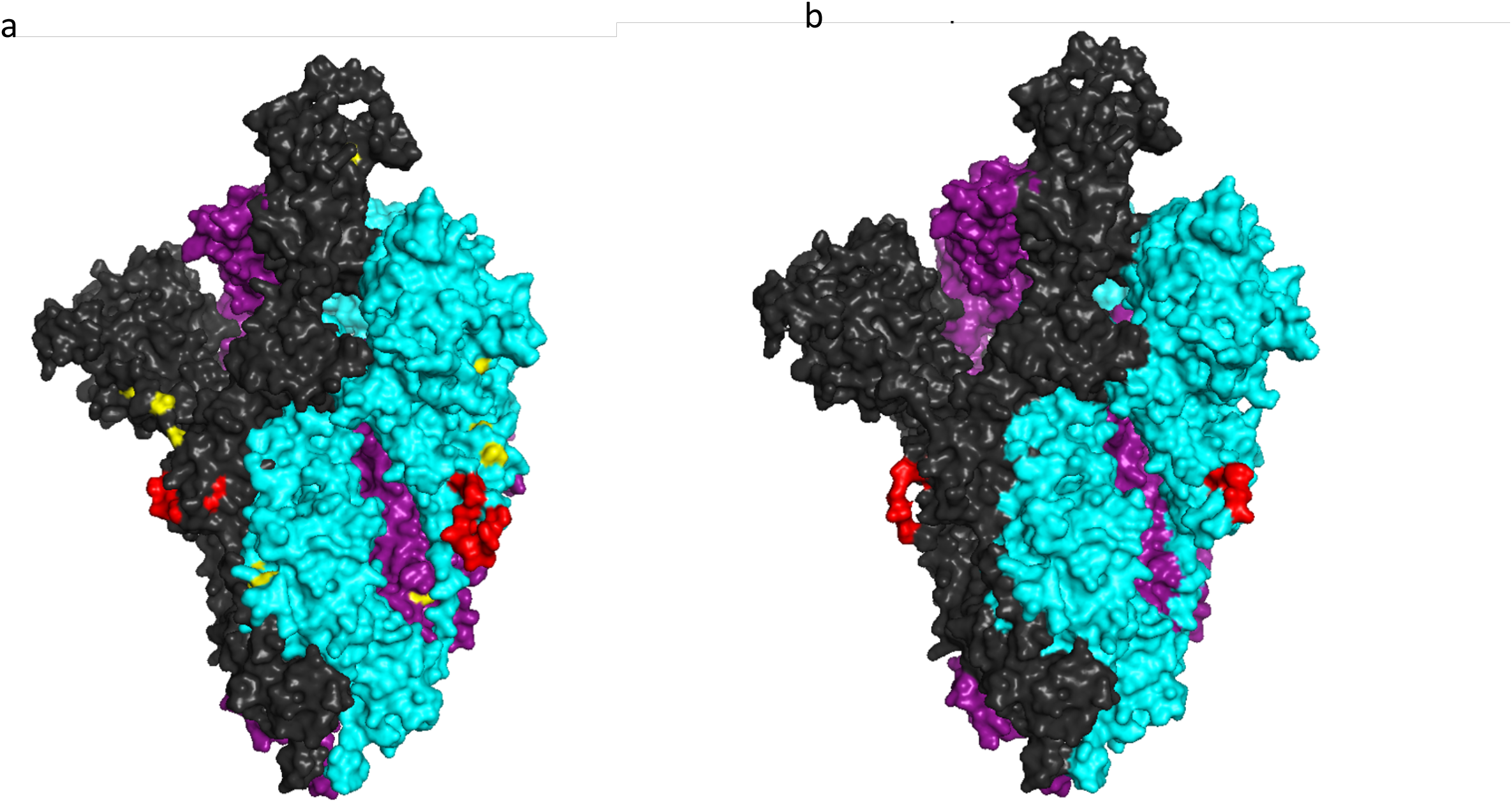
A space filled model of the wild type SARS-CoV-2 S glycoprotein in a trimeric form using the sequence of a) the native or b) spike deletant virus, in which the aa’s 679NSPRRARSV687 have been replaced with Ile. The model was built using a cryo-EM structure (6VSB.pdb) of the S glycoprotein in the prefusion form (25). Each of the monomers is coloured differently. The loop containing the furin cleavage site (or the shortened loop in the deleted version in b) is indicated in red. The positions of phosphorylation sites identified by mass spectrometry and surface located were mapped on the native structure and shown in yellow in a).

**Figure 4.**
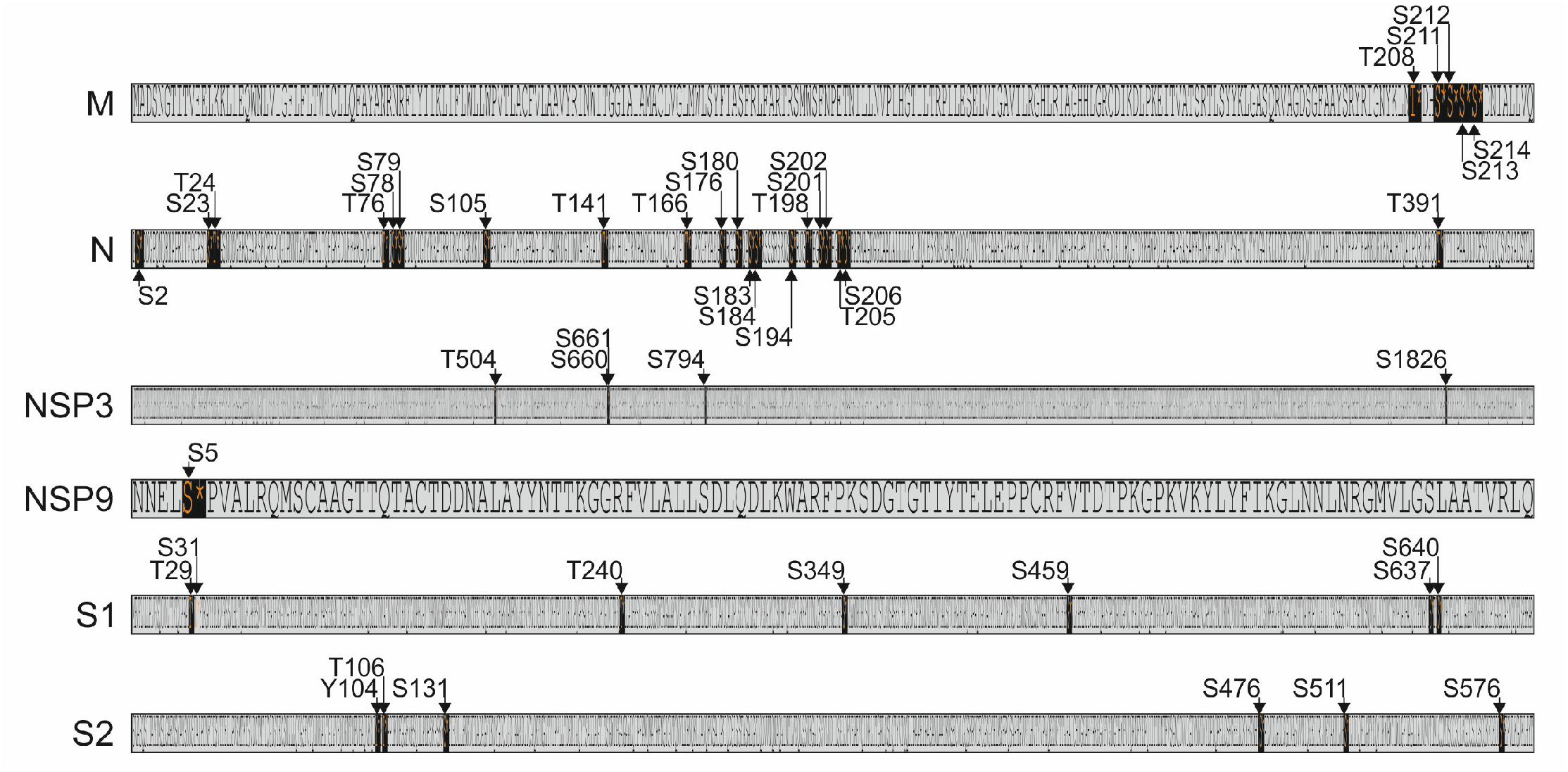
Schematic of the location of phosphorylation sites Proteins M, N, NSP3, NSP9 and S glycoprotein are shown as we have accurate phospho-site data for these proteins. For each location we indicate the aminos acid (S, T or Y) and the amino acid numbering. The S glycoprotein is shown as S1 and S2 for ease of display and to illustrate where the sites would be relative to the major cleavage site on the wild type G-glycoprotein.

**Figure 5.**
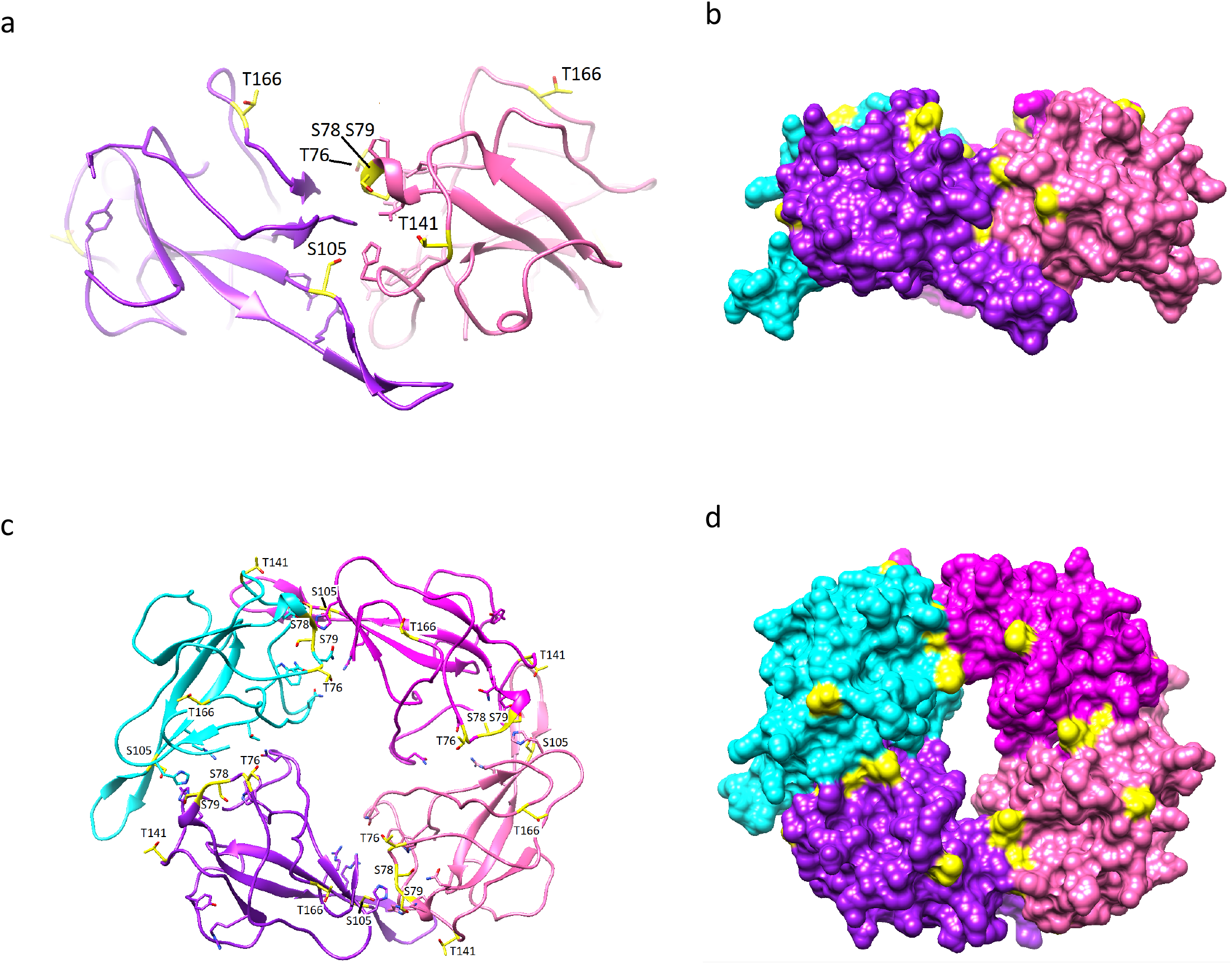
Modelling phosphorylation on the RNA binding domain of N protein. The positions of phosphorylation sites identified by mass spectrometry were mapped on the x-ray crystal structure of the N-terminal RNA binding domain of the N protein (aa residues 47 – 173) from SARS-CoV-2 (6YVO.pdb). The four monomer units in one asymmetric unit are distinctly coloured and shown as (a and b) side and (c and d) top views as ribbon (left hand figures) and space filling models (right hand figures).

## Discussion

This integrated analysis of the SARS-CoV-2 transcriptome and proteome revealed significant findings, in particular, the identification of an eight aa deletion in the SARS-CoV-2 S glycoprotein that potentially effects protein cleavage, cell tropism and infectivity. The coronavirus S glycoprotein is made as a larger precursor that must be primed by cleavage using host cell proteases to enable subsequent viral entry. For different coronaviruses, cleavage can occur at one or more sites (termed the S1/S2 and S2’ sites) depending on the amino acid consensus sequences present at each site and has major repercussions on host cell tropism and pathogenesis ^19^. The SARS-CoV S glycoprotein has a single arginine residue at the S1/S2 site (Figure 2b), that facilitates cleavage by a number of proteases including trypsin, cathepsin L and TMPRSS11D ^(32–34^ and a consensus sequence for trypsin and/or elastin cleavage at the S2’ site^(35,36^. By comparison, SARS-CoV-2 has a four amino acid insertion (_680_SPRR_683_) in the S glycoprotein that results in the generation of a furin-like cleavage site (_682_RRAR_685_) at the S1/S2 boundary that is not present in other lineage B betacoronaviruses, including SARS-CoV and the highly related bat coronavirus RaTG13^23^. Moreover, analysis of a cryo-EM structure of the S glycoprotein showed that the SPRR insertion creates a disordered solvent exposed loop^24^ that protrudes from the trimer surface (Figure 3a). It has been proposed that this region is more available to host cell proteases for processing^24^. The presence of a furin-like cleavage site at the S1/S2 boundary results in cleavage of the SARS-CoV-2 S glycoprotein before viral exit from the cell, whereas the S glycoprotein of coronaviruses such as SARS-CoV, lacking a S1/S2 furin cleavage site, exit the cells with the S glycoprotein largely uncleaved, necessitating cleavage prior to or during cell entry^20,37,38^.

The SARS-CoV-2 S glycoprotein deletion identified in this study removes the furin cleavage site, but also Arg_685_ (corresponding to SARS-CoV Arg_667_) required for cleavage of the SARS-CoV S glycoprotein. However the sequence _694_AY/T_696_ (numbering of undeleted S glycoprotein, cleavage shown by “/”; Figure 2b) is still retained and has been identified as a second potential S1/S2 protease cleavage site for SARS-CoV downstream of the basic Arg_667_ residue^23^ suggesting that either the deleted SARS-CoV-2 S glycoprotein could be cleaved at this site or only use the S2’ site for cleavage. Recently pseudoviruses expressing the SARS-CoV-2 S-glycoprotein and the corresponding S glycoprotein with the furin cleavage site removed at the S1/S2 boundary (residues _680_SPRR_683_ removed) were made and their ability to mediate entry to Vero E6 and BHK-21 cells expressing hACE2 compared^33^. Deletion of the furin cleavage site enhanced the entry of the corresponding pseudovirus into Vero E6 cells but diminished entry into hACE2 BHK-21 cells. Although the engineered deletion differed from the naturally occurring deletion described here, this observation suggests that the naturally occurring deletion (in cell culture) enhances the ability of the virus to enter Vero cells and was selected for during passage in Vero cells. This has clear implications for the use of Vero cells to propagate and grow large batches of the virus for research and especially virus batches grown for use in vaccine challenge studies. Moreover, it also raises the possibility that even virus stocks which have been carefully assayed for homogeneity could still spontaneously generate this deletion during animal challenge studies – particularly in non-human primates and perhaps especially in animals from the *Chlorocebus* genus. Thus, vaccine studies will need to carefully monitor the homogeneity of the challenge virus throughout the study period. In principle viruses carrying the Bristol deletion in the S glycoprotein may well be attenuated *in vivo* in NHPs. The early emergence of a viral population carrying the Bristol deletion within an individual challenge animal would give the false impression that an individual animal had a mild infection. Given that animal studies are required to use the smallest number of animals needed for statistical significance just one event like this could confound the study. Allied to this is the question of how such a deletion may affect therapeutic assessment of monoclonal antibodies directed against the virus. We have already instigated a programme to determine if this deletion also occurs in human clinical isolates and are currently examining the evolution of the S glycoprotein deletant virus in human cell lines to determine whether the furin cleavage site is essential for infection of human cells.

In addition to the S glycoprotein deletion we were also able to find an additional significant deletion, in high quality data recently deposited by Kim et al.^8^, in the region of the E protein. This finding reinforces the potential of the virus to incur unexpected spontaneous deletions in viral genes during passaging, that are hard to predict and detect, something researchers need to be vigilant for. A recent analysis showed that deletion of ORF8 from human clinical isolates is possible ^39^ and our data shows that novel SARS-CoV-2 viruses containing deletions or even insertions can arise naturally and successfully propagate. As the virus continues to spread and potentially comes under selective pressure from the host response in humans, vaccines and antiviral drugs in the future, vigilance will be required to detect novel rearrangements/deletions. Coronaviruses use a distinctive complex mechanism for genome replication that needs to be considered during bioinformatic analysis of their genomes relative to viruses causing other epidemics and pandemics where the principle driver of viral change would be single nucleotide polymorphisms and/or small (<3nt) insertions/deletions.

Allied to this we observed a rarer but detectable set of deletions throughout the viral mRNA repertoire with several notable deletions that give rise to uniquely identifiable peptides. This reinforces the potential for SARS-CoV-2 to naturally undergo deletions/rearrangements throughout the genome under selective pressure, something perhaps readily predictable from other coronaviruses but now confirmed directly in at least one cell line. If the plasticity of the SARS-CoV-2 genome follows the same pathway as HCoV-NL63^40^ then different isolates will emerge through recombination and this may result in repeat infection as protection will not be effective.

The detection of numerous phophorylation sites within critical viral proteins is another key resource. The phosphorylation of the N protein of multiple coronaviruses is well known^(41–45^ and the phosphorylation sites identified lie in either the N-terminal RNA binding domain or interdomain linker^46^. Mapping of the phosphorylation sites on an x-ray structure of the SARS-CoV-2 N-terminal RNA binding domain showed they were surface located (Figure 5). However, we believe this is the first report of phosphorylation of the coronavirus proteins M, nsp3 and S. These are all membrane bound proteins. M protein is critical in forming the viral particle, nsp3 is a key multifunctional component of the replication/transcription machinery and S protein is the major attachment protein respectively^47^. The phosphorylation sites identified on the S glycoprotein may be important in assembly of the trimer. Residues T29, S31, S349, T791 and S816 are all surface located, whilst T240 sits underneath a disordered loop which when phosphorylated, will add negative charge which may influence loop conformation. Residues Y789 and (perhaps to a lesser extent) T791 sit at the subunit interface and may be involved in controlling trimer assembly. S637 and S640 lie in a modelled loop but are nevertheless potentially interesting. They are close together in sequence and in the loops modelled for the compact folds these model well as hairpins. In the chain with the extended domain, the same loop models in an extended form. This would be consistent with phosphorylation at S637 and S640 forcing the hairpin apart. These are worth exploring as potential control points for spike conformational changes from compact to an extended form. Overall the identification of protein phosphorylation sites is notable but we believe that the greater utility may be that identifying numerous phophorylation sites on viral proteins will open the potential for a rational investigation of clinically licenced kinase inhibitors as antiviral drugs as has previously been observed for SARS-CoV ^48^.

Finally, this transcriptomic and proteomic dataset is a rich resource for research teams building a picture of this novel virus as it enables an accurate overview of the transcription profile and adds significant new direct observational data for the viral proteins.

## Acknowledgements

The authors would also like to thank Robin Gopal and Monika Patel from the National Infection Service at PHE Collingdale for isolation, propagation and advice on viral growth. J.A.H. would like to acknowledge the help and support of the Liverpool Malawi COVID-19 Consortium.

This work was supported by the United States Food and Drug Administration grant number HHSF223201510104C ‘Ebola Virus Disease: correlates of protection, determinants of outcome and clinical management’ amended to incorporate urgent COVID-19 studies. Awarded to J.A.H., A.D., D.A.M. and M.W.C. JAH was supported by the National Institute for Health Research Health Protection Research Unit (NIHR HPRU) in Emerging and Zoonotic Infections at University of Liverpool in partnership with Public Health England (PHE), in collaboration with Liverpool School of Tropical Medicine. The views expressed are those of the author(s) and not necessarily those of the NHS, the NIHR, the Department of Health or Public Health England.

D.A.M. and A.D.D. were supported by the BBSRC (grant BB/M02542X/1). We would also like to thank Richard Leeves and Alan Baldwin for technical help with data archiving.

## Methods

### Virus growth and assay

Vero E6 (ATCC^®^ CRL 1586^™^) cells were cultured at 37°C in Dulbecco’s modified Eagle’s medium (DMEM, Gibco^™^, ThemoFisher) supplemented with 10% fetal bovine serum (FBS), penicillin (100 units/ml) and streptomycin (100 μg/ml). All work with infectious SARS-CoV-2 strain England/2/2020 (VE6-T), isolated by Public Health England (PHE), was done inside a class III microbiological safety cabinet in a containment level 3 facility at the University of Bristol. A SARS-CoV-2 stock was produced by infecting Vero E6 cells at a multiplicity of infection (MOI) of 0.01 and incubating the cells for 72 hours. The culture supernatant was clarified by centrifugation and stored in aliquots at −80°C. The titre of the stock was determined by preparing 10-fold serial dilutions in Eagle’s minimal essential medium (MEM; Gibco^™^, ThemoFisher) supplemented with 2% FBS. Aliquots of each dilution were added to 1 × 10^4^ Vero E6 cells in the same medium in each of 12 wells of a 96-well plate. Plates were incubated at 37°C for 5-7 days and then examined for cytopathic effect. The TCID_50_ was calculated according to the method of Reed and Muench ^49^.

### RNA extraction for direct RNA sequencing

Extraction and sequencing was done as previously described ^14^. Briefly, total RNA was extracted using TRIzol^™^ reagent (#15596026, Ambion) and the RNA extracted as per the manufacturer’s recommendations but with 3 x 70% ethanol washes. After resuspension in RNAse-free water the RNA was polyA enriched and sequenced immediately using SQK-RNA002 kits and MIN106D R9 version flow cells (Oxford Nanopore Technologies).

### Data analysis, characterisation of viral transcripts

To cope with the very wide range of transcripts, and to enable grouping of transcripts into classes, our previously described ORF centric data analysis pipeline was utilised ^14^. The transcripts were mapped to the viral genome with minimap2 and the mapping data was used to identify transcription regulatory sequences (TRS), the sites where the transcript meets the poly A tail at the end of the genome and junction sites which the software refers to as splice acceptor/donor sites as it was originally used to describe spliced adenovirus transcripts. This script produces tables indicating where on the genome and how often in the data each junction occurs. Subsequently, nearby events are grouped together for simplicity of analysis. Once this is complete, the software then assigns each transcript to a “transcript group” depending on its pattern of TRS, junction sites and poly A locations and counts how many transcripts belong to each transcript group. Nanopolish^50^ was used to determine the polyA length of each sequenced transcript and subsequently the average polyA length was calculated for each transcript group.

A second in-house script determined which known features are present in each transcript group and generated pseudo transcripts based on the viral genome sequence to remove nanopore sequencing errors. The script examined each pseudo-transcript to determine what features it has, using a user-specified list of canonical features or ORFs on the viral genome (Supplementary Data 8). It also produced GFF files that allow the user to visualise the dominant transcript coding for each ORF as well as GFF files describing the whole range of transcripts coding for any given ORF. In addition, an analysis counting the final number of transcripts belonging to each translated feature was produced. The pipeline generated an ORF-centric view of the viral transcriptome – classifying transcripts according to the viral proteins coded for.

### RNA extraction for sequencing of the suspected deletion region

To prepare intracellular SARS-CoV-2 RNA, total cellular RNA containing SARS-CoV-2 RNA was extracted from the Vero E6 cells used for viral stock production using TRIzol^™^ Reagent (Invitrogen^™^, ThemoFisher) following the manufacturer’s instructions. Viral RNA was extracted from cell culture supernatants using a QIAamp Viral RNA Mini Kit (Qiagen) according to the manufacturer’s instructions. Approximately 3 kb RT-PCR products covering the S gene deletion were amplified from the viral RNA using the gene specific primers F9newF and F9newR (5’-TAAGGTTGGTGGTAATTATAATTACCTG-3’ and 5’-AAAATAGTTGGCATCATAAAGTAATGGG-3’) and a SuperScript^™^ IV One-Step RT-PCR System (Invitrogen^™^, ThemoFisher). A region spanning the deletion was sequenced using primers Wu_24_L and Wu_24_R (5’-TTGAACTTCTACATGCACCAGC-3’ and 5’-CCAGAAGTGATTGTACCCGC-3’).

### Total Proteome Analysis

Protein lysates were prepared from the Vero E6 cells used for viral stock production. The cells were harvested in 4X Laemmli buffer (BioRad) and heated to 95 °C for 15 min. A 25 μl aliquot of the sample was separated using SDS-PAGE and the gel lane cut into 20 slices. The slices were reduced (10 mM DTT, 56 °C, 30 min), alkylated (100 mM iodoacetamide, room temperature, 20 min) and digested with trypsin (0.5 μg trypsin per slice, 37 °C, overnight). This whole process was repeated with chymotryptic digestion (0.5 μg chymotrypsin per slice, 25 °C, overnight). The resulting tryptic and chymotryptic peptides were fractionated using an Ultimate 3000 nano-LC system in line with an Orbitrap Fusion Lumos mass spectrometer (Thermo Scientific). In brief, the peptides from each gel slice in 1% (vol/vol) formic acid were injected onto an Acclaim PepMap C18 nano-trap column (Thermo Scientific). After washing with 0.5% (vol/vol) acetonitrile 0.1% (vol/vol) formic acid, peptides were resolved on a 250 mm × 75 μm Acclaim PepMap C18 reverse phase analytical column (Thermo Scientific) over a 150 min organic gradient, using 7 gradient segments (1-3% solvent B over 1 min, 3-15% B over 58 min, 15-32%B over 58 min, 32-40%B over 5 min, 40-90%B over 1 min, held at 90%B for 6 min and then reduced to 1%B over 1 min) with a flow rate of 300 nl min^−1^. Solvent A was 0.1% formic acid and Solvent B was aqueous 80% acetonitrile in 0.1% formic acid. Peptides were ionized by nano-electrospray ionization at 2.2 kV using a stainless-steel emitter with an internal diameter of 30 μm (Thermo Scientific) and a capillary temperature of 250 °C.

All spectra were acquired using an Orbitrap Fusion Lumos mass spectrometer controlled by Xcalibur 4.1 software (Thermo Scientific) and operated in data-dependent acquisition mode. FTMS1 spectra were collected at a resolution of 120 000 over a scan range (m/z) of 375-1550 (for tryptic peptides) or 325-1500 (for chymotryptic peptides), with an automatic gain control (AGC) target of 4E5 and a max injection time of 50ms. Precursors were filtered according to charge state (to include charge states 2-7), with monoisotopic peak determination set to peptide and using an intensity threshold of 1E3. Previously interrogated precursors were excluded using a dynamic window (40s +/-10ppm). The MS2 precursors were isolated with a quadrupole isolation window of 0.7m/z. ITMS2 spectra were collected with an AGC target of 2E4, max injection time of 35ms and HCD collision energy of 30 %.

A targeted analysis was performed to confirm the identification of the del-spike peptide (QTQTIASQSIIA) identified by a single peptide spectral match (PSM) in the initial analysis of chymotryptic peptides. However, there were changes to the acquisition workflow. Precursors were filtered according to charge state (to include charge state 2) and previously interrogated precursors were excluded using a dynamic window (2s +/-10ppm). A targeted mass was specified with m/z 712.3759 and z=2.

### Phospho Proteome Analysis

Six 30 μl aliquots of the sample were separated by SDS-PAGE until the dye front had moved approximately 1cm into the separating gel. Each gel lane was excised as a single slice and subjected to in-gel tryptic digestion as above but using 1.5 μg trypsin per slice. The resulting peptides were subjected to TiO_2_-based phosphopeptide enrichment according to the manufacturer’s instructions (Pierce). The flow-through and washes from the TiO_2_-based enrichment were then subjected to FeNTA-based phosphopeptide enrichment, again according to the manufacturer’s instructions (Pierce). The phospho-enriched samples were evaporated to dryness and then resuspended in 1% formic acid prior to analysis by nano-LC MSMS using an Orbitrap Fusion Lumos mass spectrometer (Thermo Scientific) as above.

### Data Analysis

The raw data files were processed using Proteome Discoverer software v2.1 (Thermo Scientific) and searched against the UniProt *Chlorocebus sabaeus* database (downloaded March 2020; 19525 sequences), an in-house ‘common contaminants’ database and a custom SARS-CoV-2 protein database using the SEQUEST HT algorithm. Peptide precursor mass tolerance was set at 10 ppm, and MS/MS tolerance was set at 0.6 Da. Search criteria included oxidation of methionine (+15.995 Da), acetylation of the protein N-terminus (+42.011 Da) and methionine loss plus acetylation of the protein N-terminus (−89.03 Da) as variable modifications and carbamidomethylation of cysteine (+57.021Da) as a fixed modification. For the phospho-proteome analysis, phosphorylation of serine, threonine and tyrosine (+79.966 Da) was also included as a variable modification. Searches were performed with full tryptic or chymotryptic digestion and a maximum of 2 missed cleavages were allowed. The reverse database search option was enabled and all data was filtered to satisfy a false discovery rate (FDR) of 5%.

### Data availability

The fastq files and ThermoFisher .raw files are available on zenodo.org under the following doi’s: 10.5281/zenodo.3722580 for the fastq data, <u>10.5281/zenodo.3722604</u> for the phosphoproteomics .RAW files, <u>10.5281/zenodo.3722590</u> for the total proteome .RAW files (slices 1-10 of 20) and <u>10.5281/zenodo.3722596</u> for the total proteome .RAW files (slices 11-20 of 20).

