## Supplementary figure 1 for "Characterisation of the transcriptome and proteome of SARS-CoV-2 using direct RNA sequencing and tandem mass spectrometry reveals evidence for a cell passage induced in-frame deletion in the spike glycoprotein that removes the furin-like cleavage site"

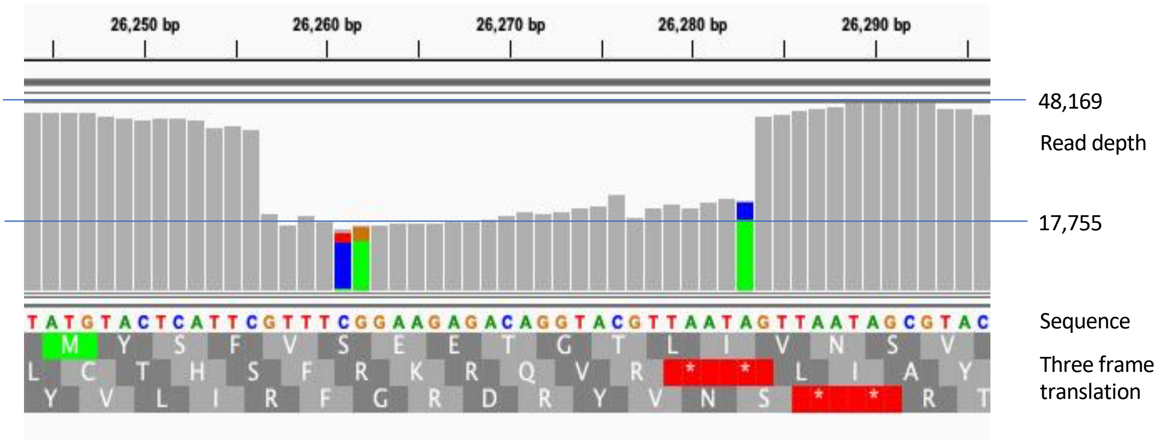


IGV viewer illustration of 27 nt deletion in the viral transcripts covering the beginning of the region coding for envelope E protein. The initiating methionine is seen in the top row of the three fram translation.
